# The generation of a Nutm1 knock-in reporter mouse line for imaging post-meiotic spermatogenesis

**DOI:** 10.1101/2022.01.04.474959

**Authors:** Maxwell Hakun, Janet Rossant, Bin Gu

**Affiliations:** Institute for Quantitative Health Science and Engineering, Michigan State University East Lansing, Michigan, United States; Department of Obstetrics, Gynecology and Reproductive Biology, Michigan State University East Lansing, Michigan, United States; Department of Biomedical Engineering, Michigan State University, East Lansing, Michigan, United States; Program in Developmental and Stem Cell Biology, Hospital for Sick Children, Toronto, ON, Canada; Department of Molecular Genetics, University of Toronto, Toronto, ON M5S 1A8, Canada; The Gairdner Foundation, Toronto, ON, Canada

## Abstract

Spermiogenesis, the post-meiotic stage of sperm development, is critical for normal male fertility. Many genetic defects and environmental assaults that affect spermiogenesis have been shown to be associated with male infertility. In addition, this later stage of spermatogenesis has been proposed to be an ideal target for male contraceptive development. The mouse is a widely used model for studying the mechanisms of spermatogenesis and spermiogenesis. However, due to the complexity and the asynchronous nature of spermatogenesis in adult testis, it is challenging to study molecular processes restricted to this specific developmental stage. It is also challenging to monitor the spermiogenesic activity in live mice, which is critical for screening for fertility-modulating interventions such as contraceptives. Here we reported the development of a Nutm1-T2A-luciferase 2(Luc2)-tandem Tomato(TdTomato) knock-in reporter mouse model that specifically labels post-meiotic spermatids. Homozygous reporter mice are healthy and fully fertile, demonstrating no interference with the normal functions of the *Nutm1* gene by the reporter. We demonstrated the visualization of post-meiotic spermatids by fluorescent imaging of the TdTomato reporter in both live and fixed testis tissues. We also demonstrated bioluminescence imaging of Nutm1 expressing cells in live mice. The Nutm1-T2A-Luc2TdTomato reporter mouse can serve as a valuable tool for studying spermiogenesis.

## Introduction

After completing meiosis, male germ cells go through profound morphological maturation and genomic reprogramming to generate functional sperm in a process called spermiogenesis^1^. Many critical processes for normal fertility, including morphological elongation, nuclear condensation, and the assembly of critical organelles such as acrosomes, proceed during spermiogenesis^1,2^. Many testes-specific genes with poorly characterized molecular functions are expressed only during spermiogenesis. A number of them have been shown to be critical for male fertility^1–3^. This makes spermiogenesis a critical stage for more mechanistic studies to better understand male infertility. Because blocking spermiogenesis processes, in general, does not interfere with the testosterone levels and the gross size and morphology of testes, it has become an intensively investigated target process for developing male contraceptives^3,4^. Indeed, two promising candidate agents for male contraceptives – JQ1 and triptonide^5,6^, both target the spermiogenesis processes. There is a clear need to be able to monitor spermiogenesis in live animals during fetal and neonatal development, puberty, aging, and under treatments of putative contraceptive drugs.

Here, we described a knock-in reporter mouse line that reports the expression of the Nuclear Protein of Testis (*Nutm1)* gene with linked fluorescent (TdTomato) and bioluminescence (Luc2) reporters : *Nutm1-T2A-Luc2TdTomato*. *Nutm1* is exclusively expressed in male germ cells after the post-meiotic mid-round spermatid stage in adult mice^7^. Genetic knockout of *Nutm1* in mice leads to infertility in male mice, likely due to defects in the protamine-histone replacement that is critical for normal spermatogenesis^7^. We demonstrated specific labeling of the post-meiotic male germ cells in live whole seminiferous tubes, as well as cryosections by TdTomato expression. The mice also presented strong testis-specific signals through bioluminescence imaging. The *Nutm1-T2A-Luc2TdTomato* mouse line can serve as a valuable tool for studying spermiogenesis in mice.

## Methods

### Designs of guide RNAs and knock-in repair donors

A guide RNA target spanning the Stop TAG codon of the *Nutm1* gene: GGTGGCCCTCTGCTTCCTACTGG was selected using the CRISPOR algorism (http://crispor.tefor.net) based on specificity scores^8^. The chemically-modified single guide RNAs (sgRNAs) with the sequence GGUGGCCCUCUGCUUCCUAC was synthesized by Synthego Inc. The repair donor for Nutm1-T2A-Luc2TdTomato reporter was designed as illustrated in Figure 1A. The T2A-Luc2TdTomato cassette coding sequences in proper orientation was flanked by long homology arms (1059bp 5’ arm and 767bp 3’arm) on each side and replaced the stop codon of the *Nutm1* gene. The homology arm sequences were PCR amplified from genomic DNA extracted from tail tissues of mice of the CD1 background. The Luc2TdTomato cassette was PCR amplified from the pcDNA3.1(+)/Luc2=TdT plasmid (Addgene, https://www.addgene.org/32904/)^9^. The T2A sequence was introduced within the upstream primer for amplifying the Luc2TdTomato cassette. The complete donor sequence of the T2A-Luc2TdTomato cassette flanked by homology arms was cloned in the EasyFusion mCherry plasmid (Addgene, https://www.addgene.org/112849/)^10^ between the EcoRI and SpeI restriction sites by multi-cassette assembly using the InFusion cloning kit (Takara Inc.). The correct sequence of the donor region was validated by Sanger sequencing.

**Figure 1:**
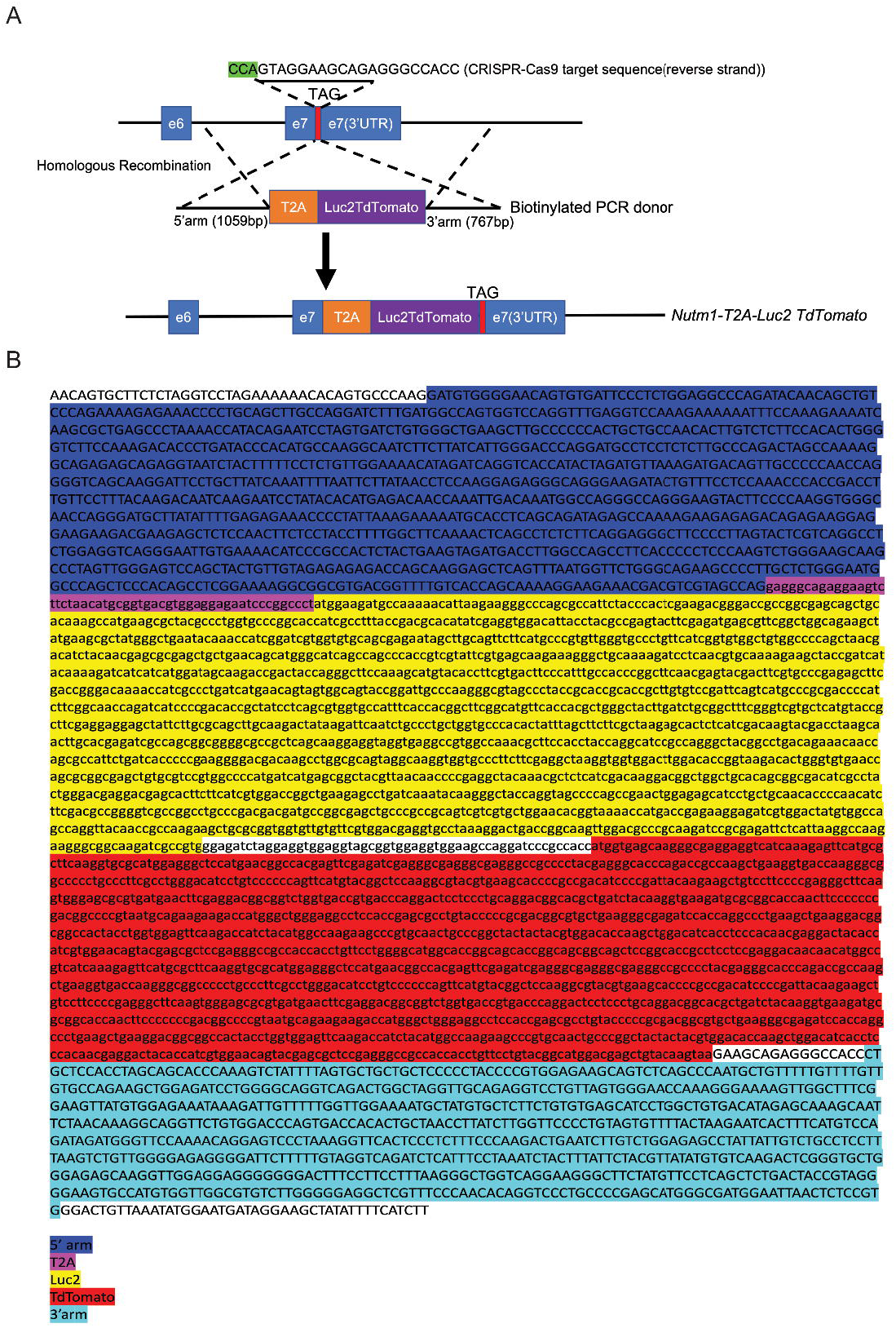
The design (A) and sequence (B) of the *Nutm1-T2A-Luc2TdTomato* allele.

### Generating KI reporter mouse lines by 2C-HR-CRISPR

The KI reporter mouse line was generated following our published protocol using 2C-HR-CRISPR on the CD1 background^10,11^. Briefly, Cas9 monomeric streptavidin (mSA) mRNA was produced by i*n vitro* transcription using the mMessage mMachine SP6 Kit (Thermo Scientific). PCR templates were generated by PCR reaction with biotinylated primers using high fidelity ClonAmp HiFi PCR mix (Takara Inc.). A mixture of Cas9-mSA mRNA (75ng/ml), sgRNA (50ng/ml), and biotinylated repair donor (20ng/ml) was microinjected into the cytoplasm of 2-cell stage mouse embryos which were transferred the same day to pseudo-pregnant females. We established founder mice with the correct insertion at high efficiency (2/10 live-born pups).

### Genotyping and genetic quality controls

Founder mice were genotyped by PCR amplification with primers spanning homology arms using the following primers: 5’ arm gtF: GAGACTGTCATAGACAGCATCCAAGAT and 5’arm gtR: tcatggctttgtgcagctgc; 3’arm gtF: ggactacaccatcgtggaacagta and 3’arm gtR: CATATTTAACAGTCCCACGGAGAG. Founder mice were out-crossed to CD1 mice to generate N1 mice. The N1 mice were genotyped by PCR. Additionally, genomic regions spanning the targeting cassette and 3’ and 5’ homology arms were Sanger-sequenced to validate correct targeting and insertion copy number was evaluated by droplet digital PCR (performed by the Centre for Applied Genomics at the Research Institute of The Hospital for Sick Children, Toronto). Heterozygous N1 mice have only one insertion copy, demonstrating single-copy insertion. An N1 founder was out-crossed five generations to wild-type CD1 mice to breed out any potential off-target mutations introduced by CRISPR-Cas9 and then bred to homozygosity at N6 generation. The mouse line was then maintained by homozygous breeding.

### Whole-mount live imaging of seminiferous tubes

Intact seminiferous tubes were dissected and isolated from the testis of adult (15 weeks old) homozygous *Nutm1-T2A-Luc2TdTomato* reporter mice following published protocols^12^. Segments of live seminiferous tubes were placed in a 50ul drop of M2 medium (Cytospring) on a 3.5cm glass-bottom culture dish (MatTek Inc.). The medium was covered with mineral oil to protect against evaporation. The sample was imaged on a Thunder Widefield Microscope (Leica) for the TdTomato fluorescent signal.

### Cryosection of testis tissues

Cryosection of testis tissues was prepared from the testis of adult (15 weeks old) homozygous *Nutm1-T2A-Luc2TdTomato* reporter mice following published protocols^13^. Sections were cut at 10μm thickness. Fresh sections were stained with Phalloidin-iFluo647 (Abcam) for F-actin and Hoechst 33258 for DNAs. The *Nutm1-T2A-Luc2TdTomato* reporter was visualized directly by the TdTomato fluorescent signal. The sample was imaged on a Thunder Widefield Microscope (Leica).

### Bioluminescence imaging

Mice were anesthetized with isofluorane gas and injected with D-luciferin (100 mg/kg, intraperitoneally). Mice were allowed to emerge from anesthesia for 15 minutes and were anesthetized again. Images were obtained using a grade 1 CCD camera cooled to −90°C using the IVIS 200 system (Xenogen Corp., Alameda, CA) with the field of view set at 10 cm in height. A photographic (gray scale) image was obtained immediately followed by the acquisition of bioluminescent images on a time scale of every 5 minutes for four timepoints. The photographic images were captured with a 0.2 to 10 second exposure time, 8 F/Stop, 4 binning, and an open filter. The bioluminescent images were captured with an exposure of 0.5 seconds, 1 F/Stop, 8 binning, and an open filter. The bioluminescent and photographic images were overlaid and analyzed using the Living Image software (Xenogen Corp.). Regions of interest were drawn around the testis signal and the total counts of photons were summed for the entire region of interest. Photon counts were further adjusted visually by normalizing the minimal (8.59E^6^) and maximal (9.64E^7^) photon counts.

### Ethical statement

All animal work was carried out following the Canadian Council on Animal Care Guidelines for Use of Animals in Research and Laboratory Animal Care under protocols approved by the Centre for Phenogenomics Animal Care Committee (20-0026H); and PROTO202000143 approved by the Michigan State University (MSU) Campus Animal Resources (CAR) and Institutional Animal Care and Use Committee (IACUC).

## Results and discussion

We set out to engineer a knock-in (KI) reporter of Nutm1 in mice (Figure 1A). To achieve strong signals for both fluorescent imaging at the single cell level and bioluminescent imaging in whole mice, we chose to use a reporter cassette consisted of a codon optimized firefly luciferase(Luc2) plus two copies of the Tomato red fluorescence protein(TdTomato) described before^9^. To avoid any interference to the functions of the endogenous NUTM1 protein by this large reporter cassette, we separated the coding sequence of *Nutm1* and the reporter cassette with a thosea asigna virus 2A (T2A) self-cleavage sequence. The T2A-Luc2TdTomato cassette was inserted into the endogenous *Nutm1* locus using 2C-HR-CRISPR, replacing its stop codon^10^. We performed extensive quality control to confirm single-copy insertion of the fusion reporter with the correct sequence, as detailed in the Methods section and Figure 1B and our published protocol^10,11^. The homozygous mice are healthy and fertile. The mouse line was maintained in the homozygous state after outcrossing for five generations to ensure no carry-over of any possible off-target alterations.

To evaluate the expression pattern of the *Nutm1-T2A-Luc2TdTomato* in the testis, we performed fluorescent imaging. As shown in Figure 2A, in live whole mount seminiferous tubes, red fluorescent signals are seen in cells close to the inner channel of the seminiferous tube, consistent with the anatomical position of post-meiotic germ cells. These signals were not observed in wild-type control mice, demonstrating the specificity of the reporter signals. In addition, we imaged cryosections of seminiferous tubes. DNA (stained by Hoechst 33258) and F-actin (stained by phalloidin-iFluo647) stains were used to visualize the morphological traits of male germ cells of different developmental stages in the seminiferous tubes. Again, compared to wildtype control samples, specific red fluorescence signals were observed in post-meiotic germ cells, consistent with previous reports by immunofluorescence (Figure 2B). Thus, the *Nutm1-T2A-Luc2TdTomato* reporter specifically marks the post-meiotic germ cells in the male germline in adult mice.

**Figure 2:**
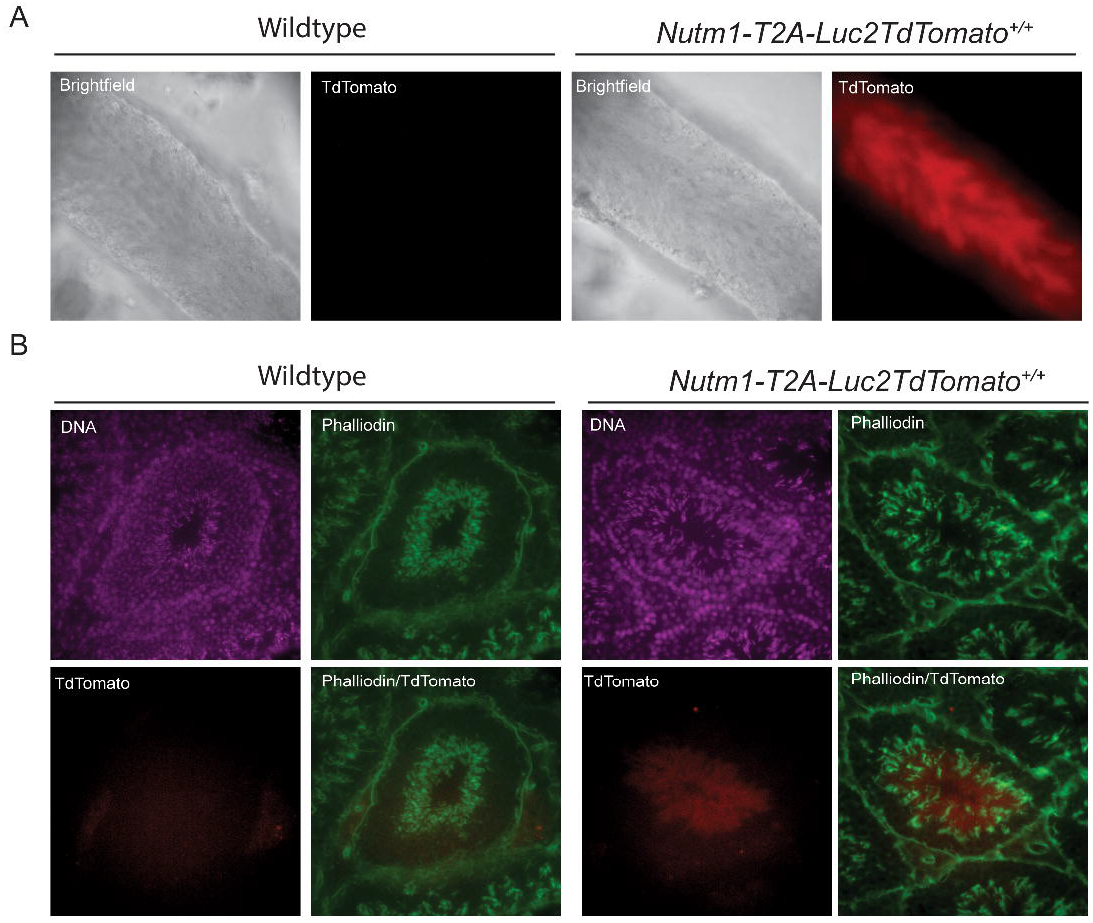
Expression pattern of the *Nutm1-T2A-Luc2TdTomato* in seminiferous tubes. A. Whole mount images of live seminiferous tubes from a wild-type control and a *Nutm1-T2A-Luc2TdTomato*^+/+^ seminiferous tube segment. B. Fluorescence images of cryosections of a wild-type control and a *Nutm1-T2A-Luc2TdTomato*^+/+^ seminiferous tube.

We then evaluated the expression pattern of *Nutm1-T2A-Luc2TdTomato* in living mice by bioluminescence imaging. As shown in Figure 2A, in *Nutm1-T2A-Luc2TdTomato*^+/+^ males, specific signals in the testis region were observed. The signal strength is slightly lower in a 35-week-old male for which the fertility is declining than a highly fertile 15-week-old male, suggesting this model might be used to monitor age-dependent fertility change in the future after more detailed characterization. No signal was observed in a *Nutm1-T2A-Luc2TdTomato*^+/+^ female, nor in a wildtype control male, demonstrating the specificity of the signal. Also, no signal was observed in any anatomical location other than the testis region. Although there are anecdotal reports that suggest the expression of *Nutm1* in non-testis tissues at the RNA level^14^, it seems that robust expression that detectable by bioluminescence imaging only existed in the testis. Thus, the *Nutm1-T2A-Luc2TdTomato* reporter can specifically report the post-meiotic germ cells in living mice by bioluminescent imaging.

In mice, the first wave of spermatogenesis follows a unique temporal pattern. The post-meiotic spermatids start to emerge around 18-20 days of age and reach the adult levels at around 25 days of age^7^. Thus, we conducted a time series of biolumiscence imaging in a young *Nutm1-T2A-Luc2TdTomato*^+/+^ male. As shown in Figure 3B, very little signal was observed in the mouse at 12 days. The signal started to increase at 20 days and reached a high level at 25 days, consistent with the known pattern of timing of spermiogenesis. However, the biluminescence signal at day 12 is non-zero, which may suggest some level of expression of the *Nutm1* gene in early spermatogenesis before meiosis. More detailed investigation is needed to further characterize the *Nutm1* expression at early spermatogenesis stages.

**Figure 3:**
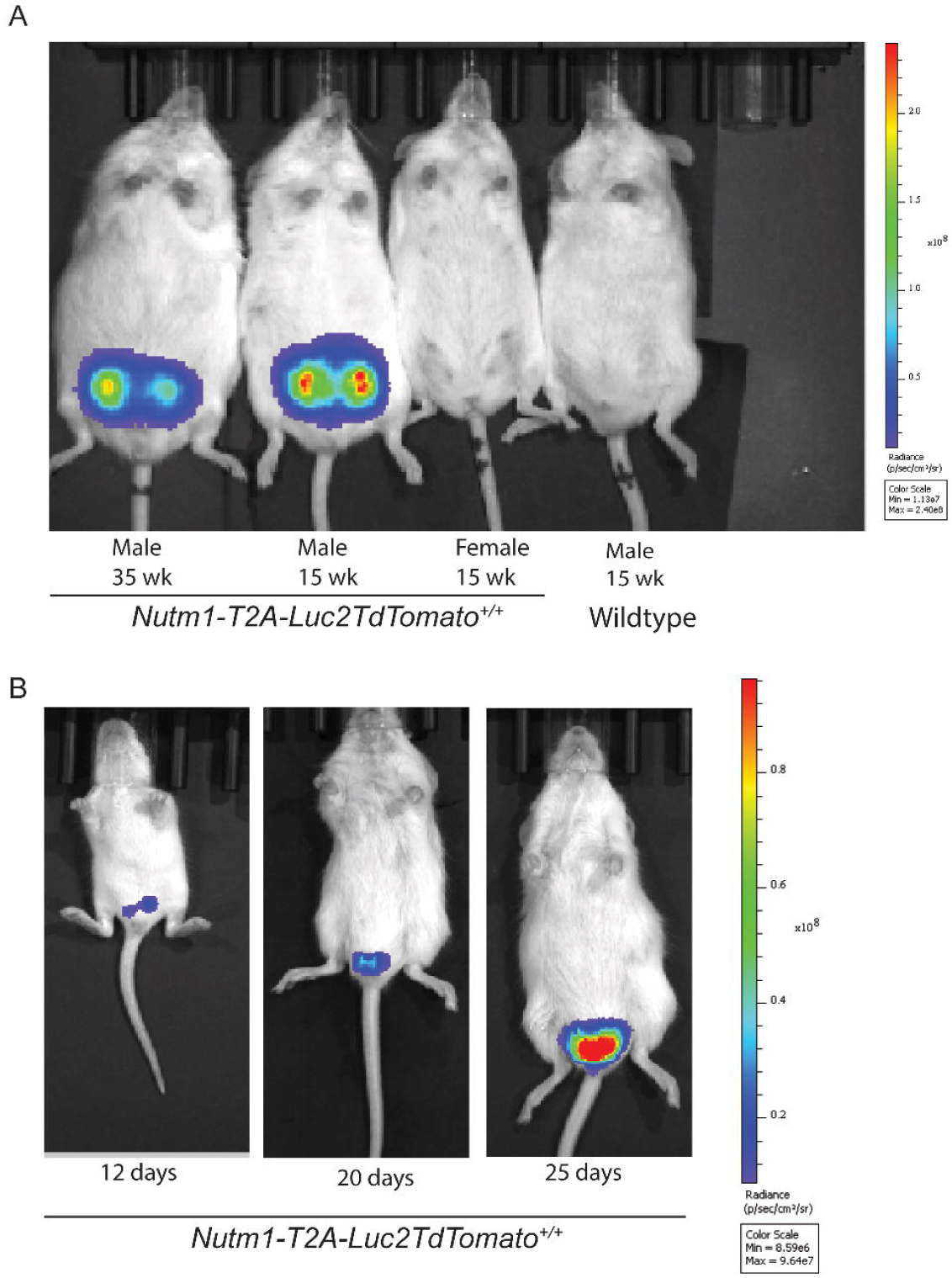
Bioluminescent imaging of the *Nutm1-T2A-Luc2TdTomato* reporter mice. A. BLI images of *Nutm1-T2A-Luc2TdTomato* in adult mice. B. A. BLI images of *Nutm1-T2A-Luc2TdTomato* in young mice.

In summary, we report a knock-in reporter mouse line that allows the imaging of post-meiotic male germ cells in vivo. The mouse line can serve as a valuable tool for studies on male fertility that require dynamic information, including the mechanisms of temporal coordination of spermatogenesis waves, impacts of aging on spermatogenesis and development of reversible contraceptives.

### Broader applications

Although *Nutm1* specifically expresses and functions in the male germline under physiological conditions, it is also a recurrent fusion partner for a large group of poorly understood cancers named *Nutm1-*rearranged neoplasms^15^. Our *Nutm1-T2A-Luc2TdTomato* reporter line may contribute to future research into these cancers and have broader applications outside reproductive biology.

## Acknowledgement

This work was funded by CIHR (FDN-143334) to J.R and Bin Gu’s start-up Funding from Michigan State University. The authors acknowledge technical support from the Model Production Core staff led by M. Gertsenstein at the Centre for Phenogenomics; The Luc2-TdTomato construct is a kind gift from Dr. Christopher Contag.

## Author contributions

B.G., conceived the study. B.G. designed and produced the *Nutm1-T2A-Luc2TdTomato* reporter mouse line. B.G and M.H designed, carried out and analyzed all experiments. J.R. and B.G. provided supervision and funding for the study. B.G wrote the manuscript and all authors reviewed and approved the manuscript.

## References

1 O’Donnell, L. Mechanisms of spermiogenesis and spermiation and how they are disturbed. Spermatogenesis 4, e979623, doi:10.4161/21565562.2014.979623 (2014).

2 Azhar, M. et al. Towards Post-Meiotic Sperm Production: Genetic Insight into Human Infertility from Mouse Models. Int J Biol Sci 17, 2487–2503, doi:10.7150/ijbs.60384 (2021).

3 Yan, W. Male infertility caused by spermiogenic defects: lessons from gene knockouts. Mol Cell Endocrinol 306, 24–32, doi:10.1016/j.mce.2009.03.003 (2009).

4 Chen, S. R. et al. The control of male fertility by spermatid-specific factors: searching for contraceptive targets from spermatozoon’s head to tail. Cell Death Dis 7, e2472, doi:10.1038/cddis.2016.344 (2016).

5 Matzuk, M. M. et al. Small-molecule inhibition of BRDT for male contraception. Cell 150, 673–684, doi:10.1016/j.cell.2012.06.045 (2012).

6 Chang, Z. et al. Triptonide is a reversible non-hormonal male contraceptive agent in mice and non-human primates. Nat Commun 12, 1253, doi:10.1038/s41467-021-21517-5 (2021).

7 Shiota, H. et al. Nut Directs p300-Dependent, Genome-Wide H4 Hyperacetylation in Male Germ Cells. Cell Rep 24, 3477–3487 e3476, doi:10.1016/j.celrep.2018.08.069 (2018).

8 Concordet, J. P. & Haeussler, M. CRISPOR: intuitive guide selection for CRISPR/Cas9 genome editing experiments and screens. Nucleic Acids Res 46, W242–W245, doi:10.1093/nar/gky354 (2018).

9 Patel, M. R. et al. Longitudinal, noninvasive imaging of T-cell effector function and proliferation in living subjects. Cancer Res 70, 10141–10149, doi:10.1158/0008-5472.CAN-10-1843 (2010).

10 Gu, B., Posfai, E. & Rossant, J. Efficient generation of targeted large insertions by microinjection into two-cell-stage mouse embryos. Nat Biotechnol 36, 632–637, doi:10.1038/nbt.4166 (2018).

11 Gu, B., Posfai, E., Gertsenstein, M. & Rossant, J. Efficient Generation of Large-Fragment Knock-In Mouse Models Using 2-Cell (2C)-Homologous Recombination (HR)-CRISPR. Curr Protoc Mouse Biol 10, e67, doi:10.1002/cpmo.67 (2020).

12 Yao, C. et al. Seminiferous tubule molecular imaging for evaluation of male fertility: Seeing is believing. Tissue Cell 58, 24–32, doi:10.1016/j.tice.2019.04.003 (2019).

13 Sato, T. et al. Testis tissue explantation cures spermatogenic failure in c-Kit ligand mutant mice. Proc Natl Acad Sci U S A 109, 16934–16938, doi:10.1073/pnas.1211845109 (2012).

14 Eagen, K. P. & French, C. A. Supercharging BRD4 with NUT in carcinoma. Oncogene, doi:10.1038/s41388-020-01625-0 (2021).

15 Luo, W. et al. NUTM1-Rearranged Neoplasms-A Heterogeneous Group of Primitive Tumors with Expanding Spectrum of Histology and Molecular Alterations-An Updated Review. Curr Oncol 28, 4485–4503, doi:10.3390/curroncol28060381 (2021).

